# MiMIC analysis reveals an isoform specific role for *Drosophila* Musashi in follicle stem cell maintenance and escort cell function

**DOI:** 10.1101/2020.08.25.265769

**Authors:** Nicole A. Siddall, Franca Casagranda, Timothy M. Johanson, Nicole Dominado, James Heaney, Jessie M. Sutherland, Eileen A. McLaughlin, Gary R. Hime

## Abstract

The *Drosophila* ovary is regenerated from germline and somatic stem cell populations that have provided fundamental conceptual understanding on how adult stem cells are regulated within their niches. Recent ovarian transcriptomic studies have failed to identify mRNAs that are specific to follicle stem cells (FSCs), suggesting that their fate may be regulated post-transcriptionally. We have identified that the RNA-binding protein, Musashi (Msi) is required for maintaining the stem cell state of FSCs. Loss of *msi* function results in stem cell loss, not due to cell death, but mutant FSCs upregulate Lamin C, indicating a change in differentiation state. In *msi* mutant ovaries, Lamin C upregulation was also observed in posterior escort cells and mutant somatic cells within regions 2/3 were dysfunctional, as evidenced by the presence of germline cyst collisions and fused egg chambers. The *msi* locus produces two classes of mRNAs (long and short). We show that FSC maintenance and escort cell function specifically requires the long transcripts, thus providing the first evidence of isoform-specific regulation in a population of *Drosophila* epithelial cells. We further demonstrate that although male germline stem cells have previously been shown to require Msi function to prevent differentiation this is not the case for female germline stem cells, indicating that these similar stem cell types have different requirements for Msi, in addition to the differential use of Msi isoforms between soma and germline.

## Introduction

Adult stem cells possess both the ability to self-renew and give rise to differentiated cells in order to maintain tissue homeostasis in multicellular organisms (1). Generally, stem cell self-renewal must occur through one of two mechanisms; either by asymmetric division, or at the population level by population asymmetry (2). Most adult stem cells reside in specialized environments (niches) where they receive signals that regulate their behaviour (reviewed in (3)). The coupling of extracellular signals with intrinsic activation of transcriptional networks governs a stem cell’s ability to self-renew or differentiate. Additional layers of genetic complexity reside in the form of differential transcript isoform usage, and a range of other post-transcriptional mechanisms, to ensure the proper protein diversity necessary for the regulation of stem cell homeostasis and differentiation (4, 5).

The adult *Drosophila* ovary provides an excellent model system for the study of stem cell behaviour and organ morphogenesis. The ovary is comprised of approximately 16-18 individual ovarioles which are sequential chains of egg chambers, the most mature chambers being found furthest from the anterior germarium (6). The germarium is divided into 3 regions (Fig. 1A). Germline stem cells (GSCs), in region 1, reside within a niche of somatic cap cells (CC), terminal filament cells (TF) and escort cells (ECs). GSCs divide asymmetrically to produce daughter cystoblasts (CBs). CBs divide four times synchronously with incomplete cytokinesis to form mitotic cysts of 2, 4, 8 and 16 cell cysts (6). ECs, also known as inner germarial sheath (IGS) cells, engulf cystoblasts and germline cysts in region 1 and 2a (7). Follicle cells (FCs), derived from follicle stem cells (FSCs), surround cysts in region 2b and stage 1 egg chambers are formed in region 3 (6). Somatic ECs are typically quiescent and support the developing germline, with tight association of ECs with the germline being necessary for germline development (7–10). ECs differ in their shape, size and ability to associate with germ cells depending on their position within the germarium suggesting that ECs are functionally diverse (7). Recent single-cell analysis of the adult ovary has uncovered distinct subpopulations of escort cells which physically interact with different developmental stages of GSC progeny from region 1 to the posterior region of 2a (11–14). Importantly, the different EC populations have been shown to have distinct functions in the regulation of GSC lineage development (11, 12).

**Figure 1.**
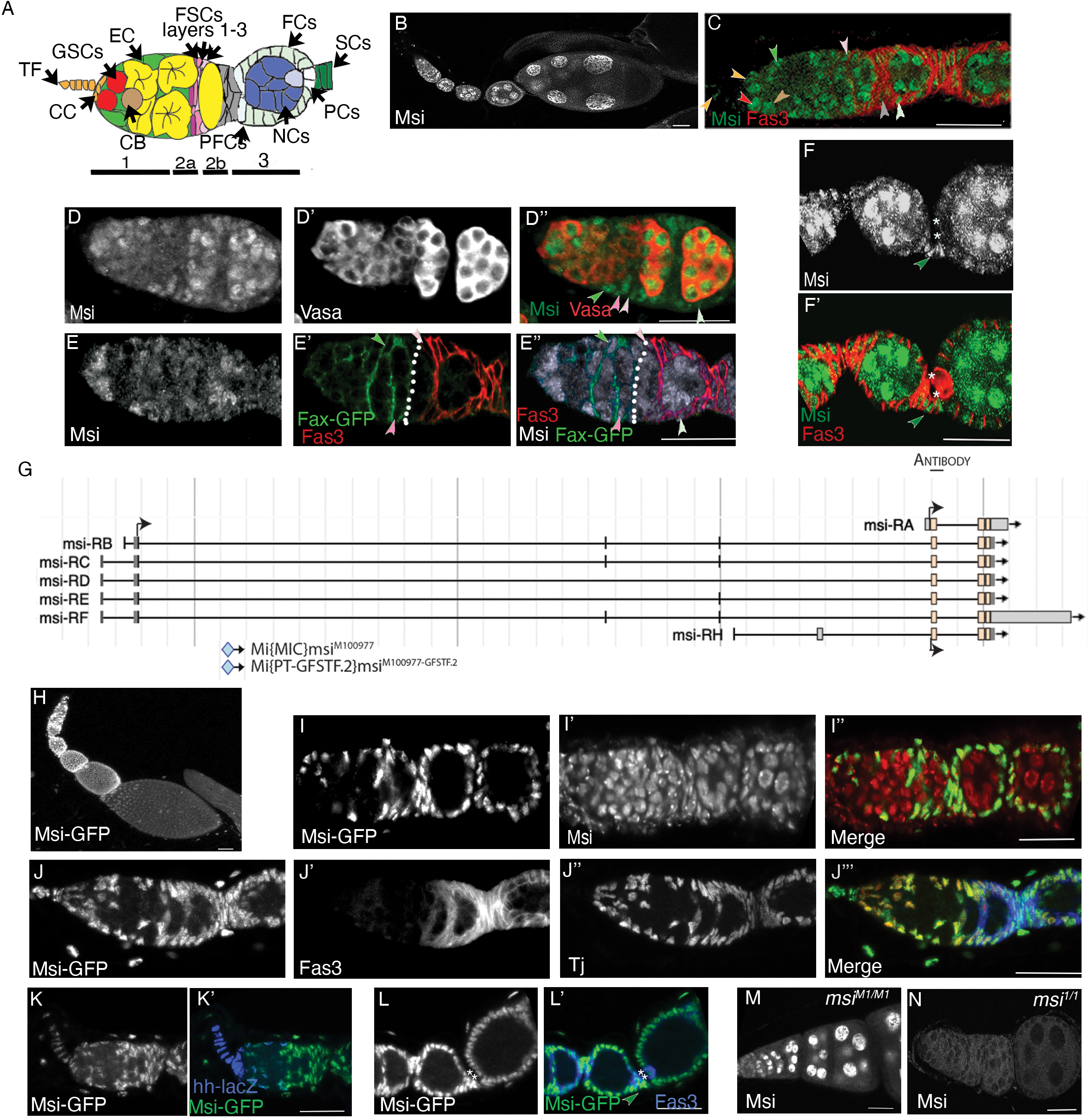
Differential expression of Msi isoforms in the adult ovary. (A) Cartoon depiction of the beginning stages of *Drosophila* ovary development showing the cell types, including terminal filament cells (TF, orange) and cap cells (CC, orange), escort cells (EC, green), germline stem cells (GSC, red), a cystoblast (CB, brown), germ cell cysts (yellow), follicle stem cells layers 1-3 (FSCs, layer 1 light pink, layer 2 medium pink, layer 3 mauve), pre-follicle cells (PFC, grey), polar cells (PC, white), nurse cells (NC, dark blue), differentiated follicle cells (FCs, light green) and stalk cells (SCs, dark green). (B-F’) Confocal micrographs of Msi antibody expression in wild-type ovaries. (B) Low magnification micrograph of ovary showing Msi expression in the germline and somatic cells of the ovary. (C) Higher magnification micrograph of ovariole labelled with Msi (green) and Fas3 (red). Msi expression was observed in GSCs (red arrowhead), CBs (brown arrowhead), ECs (green arrowhead) and in somatic TFs and CCs (orange arrowheads). In this ovariole, a Msi-expressing layer 1 FSC is labelled (light pink arrowhead). Msi expression was observed in PFCs (grey arrowhead) and differentiated FCs (light green arrowhead). A reduction of visible Msi expression in 4-8 cell germline cysts was consistently observed. (D-D”) High magnification image of ovariole labelled with Vasa (red in D”) and Msi (green in D”). Msi-positive FSCs in layer 1 (light pink arrowhead) and layer 2 (medium pink arrowhead) are labelled. The green arrowhead labels a Msi-positive posterior EC, while the light green arrowhead points to a Msi-expressing differentiated FC. (E-E”) A confocal micrograph of an ovariole labelled with Msi (grey in E”), Fax-GFP (marking the membranes of ECs; green in E”) and Fas3 (red in E”) showing a Msi-expressing layer 1 FSC (light pink arrowhead) and layer 2 FSC (dark pink arrowhead) at the 2a/b boundary (white dotted line). (F-F”). Single-plane confocal image showing Msi expression (green in F’) in later stage egg chambers also labelled with Fas3 (red). Msi expression was observed in nurse cells, mature follicle cells and stalk cells (dark green arrowhead). * Denotes the polar cells, where Msi expression was barely detectable by immunofluorescence. (G) Image modified from Flybase (J-Browse) depicting 7 *msi* transcripts. Coding start sites are labelled (black arrow), but for representation only one arrow shows that isoforms B,C,D,E and F have the same coding start site. The genomic region of the peptide sequence used to generate the Msi antibody is labelled and the insertion site of Mi{MIC}msi^M100977^, referred to as the *msi^M1^* allele in our paper, is shown. The Msi-GFP protein trap, also shown in this schematic, had been generated from the Mi{MIC}msi^M100977^ insertion. (H-N) Confocal micrographs mapping Msi-GFP expression in early oogenesis. (H) Low magnification projected image of Msi-GFP expression in the ovary shows GFP in the somatic cells. (I-I”) Confocal micrograph of ovariole labelled with the Msi antibody (red in I”) and Msi-GFP (green in I”) reveals that Msi-GFP is expressed in somatic cells but not the germline of the ovary. (J-J”’) Projection of 3 single-plane confocal images from a z-stack shows Msi-GFP expression (green in J”’) completely overlaps with TJ-positive somatic cells (red in J”’) in the germarium and early-stage egg chambers. (K-K”) Msi-GFP expression (green in K’) overlaps with *hh*-lacZ expression (blue in K’) in TFs and CCs of ovarioles. (L-L’) Msi-GFP (green in L’) is expressed in SCs (dark green arrowhead) and PCs (asterisks). (M) Single-plane confocal image of ovariole dissected from a *msi^M1/M1^* homozygous adult shows Msi antibody expression in germ cells, but not somatic cells. (N) No Msi antibody expression was detected in ovarioles dissected from a *msi^1/1^* null homozygous adult. Scale bars, 20μm.

Follicle stem cells (FSCs) reside in region 2a/2b of the germarium (15), the exact number of which remains controversial. Early studies proposed that 2-3 somatic FSCs divide to give rise to a daughter stem cell and follicle precursor cell (FPCs). FPCs then differentiate into posterior follicle cell (FC) types, including polar, stalk and main body epithelial cells (15). A more recent study which utilised a novel clonal marking system proposed that a population of approximately 14 FSCs exist in three layers within region 2a/2b (16). This latter study also suggested that it’s the FSCs in the most posterior layer (layer 1) that give rise to the follicle cell lineage, while cells within layer 3 have the capacity to differentiate into more anterior escort cells. The complexities of identifying a distinct population of FSCs cannot be understated, highlighted by the inability of single-cell RNA sequencing projects to define a clear FSC population (13). Slaidina and colleagues (2021) suggested that maybe this was because the FSC gene expression profile was likely to be similar to pre-follicle cells. In a separate study, Shi and colleagues (2021), in their single cell RNA-sequencing analysis of somatic escort and follicle cells, failed to identify any cell clusters displaying increased or decreased mitotic activity and suggested that mitotic activity between FSCs and ECs are not significantly different. The lack of clarity in the identification and function of FSCs and surrounding somatic cells highlights the need for the identification of individual genes that affect differentiation of these cell types.

Numerous studies report the importance of RNA-binding proteins (RBPs) as essential regulators of stem cell maintenance across both vertebrates and invertebrates. In *Drosophila*, RBPs have been shown to play a key role in regulation of the GSC lineage in both males and females (17–21). We previously demonstrated an intrinsic requirement for the RBP Musashi (Msi) in GSC maintenance in the *Drosophila* testis (18). In the *Drosophila* midgut, overexpression of Msi has been shown to promote intestinal stem cell proliferation in radiation-induced damaged intestines (22). Its vertebrate orthologues, Msi-1 and Msi-2, have both been described as having important functions in the regulation, proliferation and maintenance of neural, gastrointestinal, mammary, stomach and hematopoietic stem cells (23–30) and play key roles in vertebrate spermatogenesis (31, 32) and folliculogenesis (33).

The recent discovery of isoform specific functions for vertebrate Msi-1 or Msi-2 in processes such as tumor progression (34, 35) has added additional complexities to understanding the role Musashi proteins play in developmental processes. According to Flybase (FB2022_02), *Drosophila* Msi has 7 annotated transcripts and 7 polypeptides, 5 of which are unique (35). To date there has been no research that has described any isoform specificity in relation to *Drosophila* Msi function in developmental processes. By using flies carrying a Minos-Mediated Integration Cassette (MiMIC) insertion in only a subset of Msi isoforms, and a Msi protein-trap carrying GFP (recombineered from the MiMIC transposon), we describe how we have been able to uncover expression differences in Msi isoforms in the soma and germline of the *Drosophila* ovary. Additionally, we demonstrate an isoform specific requirement for Msi in the regulation of FSC maintenance and in the function of a subpopulation of ECs to support germline cyst progression in early oogenesis. In contrast, there seems to be no requirement for an alternative Msi isoform in the regulation of female GSCs, despite this same isoform being intrinsically required for maintaining spermatogonial GSC fate (18), revealing sex-specific functions for Msi. Importantly, our study is the first to identify a functional requirement for alternative Msi isoforms in different *Drosophila* stem cell populations.

## Results

We previously demonstrated an intrinsic requirement for Msi in male GSCs to maintain GSC identity (18). To investigate whether Msi is similarly required in the stem cell populations of the ovary, we first mapped the expression pattern of Msi in cells at the early stage of oogenesis. Analysis of Msi protein distribution in the ovary using a polyclonal antibody specific to Msi (Fig. 1N, Supplementary Figure 1A) (18, 36) showed Msi expression in ovarian germline and somatic cells (Fig. 1B-F’). In the germarium, Msi expression was observed in GSCs and differentiated germ cells (Fig. 1B-D). Similar to our observations in male germ cells (18), a reduction of Msi protein expression levels was consistently observed by immunofluorescence in 4-8 cell germline cysts within the germarium (Fig. 1C-E”). Msi expression persisted in nurse cells in later stage egg chambers but was absent from the oocyte (Fig. 1B). In somatic cells, Msi expression was observed in TFs, CCs, ECs and FSCs (Fig. 1C-E”) and in differentiated FCs and stalk cells (Fig. 1C-F’), but expression was barely detectable by immunofluorescence in polar cells (Fig. 1F-F’). Fasciclin 3 (Fas 3) expression is maintained at high levels in polar cells despite being reduced in mature epithelial cells, making it a suitable polar cell marker (Fig. 1F), while FSCs were identified as somatic cells located at the region 2a/2b junction, with the most posterior layer 1 cells located at the Fas 3 boundary where membranes of cells are labelled with low levels of Fas 3 (Fig. 1C-E) (37).

Protein traps serve as an additional means to analyse expression patterns of proteins. *Drosophila msi* has 7 annotated transcripts and 7 polypeptides, 5 of which are unique (Fig. 1G) (35). A Msi-GFSTF (Msi-GFP) protein trap line, generated by recombination mediated cassette exchange (RMCE) from a Mi(38)msi^M100977^ insertion in a coding intron of 5 of the 7 transcripts, was available from Bloomington Stock Centre. Msi-GFP would recapitulate expression of longer Msi isoforms, but not shorter isoforms Msi-PA and Msi-PH (Fig. 1G). Conversely, the Msi polyclonal antibody generated against amino acids 1-210 of Msi-PA would detect expression of all Msi isoforms (36) (Fig. 1G, Supplementary Figure 2).

Analysis of Msi-GFP expression in the ovary yielded surprising results with the discovery of GFP expression in only the somatic cells of the ovary (Fig. 1H-L’). Msi-GFP expression overlapped with Msi antibody expression in somatic cells (Fig. 1-I”) but was absent in the germline. Msi-GFP expression was observed in all somatic cells labelled with anti-Traffic Jam (TJ) (39) which marks CCs, ECs, FSCs and progeny (Fig. 1 J-J’’’). Co-expression of GFP and *hh-lacZ* highlighted Msi expression in the TF cells (Fig. 1K-K’). GFP expression was detected in stalk cells and polar cells (Fig. 1L-L’) despite expression being undetectable in polar cells using the Msi antibody. This result is likely due to the augmented ability to detect GFP by indirect immunofluorescence. Interestingly, these results reveal a divergence of expression of short Msi isoform/s (Msi-PA and/or Msi-PH) and the longer Msi isoforms in the ovary.

To understand the nature of the Mi{MIC}msi^M100977^ insertion (which from hereon will be referred to as *msi^M1^*) and determine its effect on Msi expression in the ovary, we tested whether Msi protein expression could be detected in homozygous *msi^M1^* flies. MiMIC insertions, when inserted in coding introns in the same orientation as the transcript, should function as gene traps if they carry a splice acceptor site followed by stop codons in all three reading frames in addition to a polyadenylation signal (40). Such gene traps can disrupt gene function. In most cases the RMCE event also rescues protein function. *msi^M1^* homozygotes are viable as adults but the presence of the MiMIC insertion is expected to disrupt expression and function of the longer Msi isoforms (Fig. 1G). While Msi antibody expression persisted in germ cells in *msi^M1/M1^* mutant ovaries, no expression was detected in somatic cells by immunofluorescence (Fig. 1M). These results demonstrate knockdown of Msi in the somatic cells of *msi^M1/M1^* mutant ovaries, but not the germline, and is consistent with expression analysis using Msi-GFP and the Msi polyclonal antibody. Together, our results suggest a divergence of Msi isoform usage in the germline and somatic cells of the ovary. Msi expression divergence was also observed in the testis. Msi antibody expression in the adult testis has been observed in somatic hub cells, cyst progenitor cells, in germline stem cells and throughout the germline (18). In adult *msi^M1/M1^* mutant testes Msi antibody expression persisted in the germline and early cyst cells but was absent from hub cells and more mature cyst cells, suggesting that testis germ cells express Msi-RA/RH but not the longer Msi isoforms (Supplementary Fig. 3A). This correlated with Msi-GFP expression, which was observed in hub, cyst progenitor and cyst cells, but was absent from the germline (Supplementary Fig. 3B).

We next sought to investigate whether Msi is functionally required for early ovary morphogenesis. Homozygous *msi* null mutants (*msi^1/1^*) are viable into early adulthood, albeit they are less fit than their heterozygous and wild-type counterparts. Their ability to survive a few days into adulthood, however, provides an opportunity to undertake morphological analysis by immunofluorescence in null mutant adult tissue. The *msi^1^* mutation was originally generated by imprecise excision of a P{LacZ} enhancer trap element near the 5’ region of the short *msi-RA* isoform (41). Msi antibody expression was not detected in ovaries dissected from *msi^1/1^* null mutants (Fig. 1N) confirming knockdown of all Msi isoforms. Germaria dissected from *msi^1/1^* 2-3 day old adults were swollen in appearance and contained excess Vasa-labelled germ cells (Fig. 2A-A’). Using the germ-cell marker Vasa coupled with the fusome marker 1B1 to identify germline cysts, we counted significantly more 4-16 cell germarial cysts in *msi^1/1^* mutant ovarioles compared to control *msi^1/+^* ovaries (Fig. 2C). A similar result was observed in ovaries from flies with a heteroallelic combination of the *msi^2^* hypomorphic allele (41) (Supplementary Fig. 1A) and the *msi^1^* null allele (Fig. 2C).

**Figure 2.**
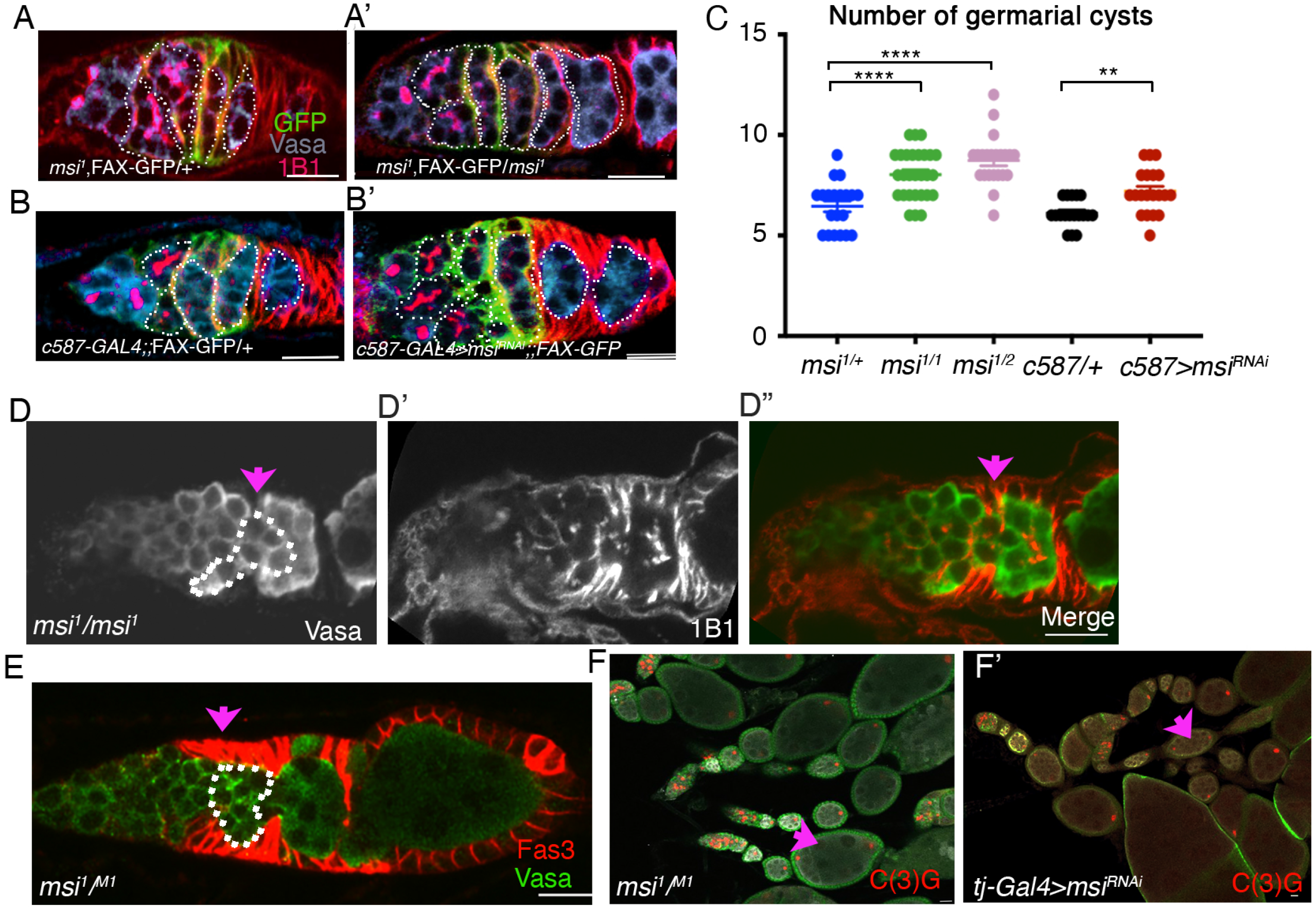
Loss of Msi from somatic cells results in an increase in the number of germarial cysts, germline cyst collisions in region 2a/2b and fused egg chambers. (A-B’) Confocal micrographs of control (A and B), *msi* null (A’) and *C587-Gal4>UAS-msiRNAi* (B’) germaria. Germline cysts (dotted outlines) were counted in germaria labelled with Vasa (germ cells), 1B1 (fusomes showing connections between germ cells), and FAX-GFP, a marker of escort cell cytoplasm. (C) Scatterplot of the average number of 4-16 cell germline cysts in regions 1 to 3 of germaria (± SEM) dissected from heterozygote *msi^1/+^* (blue; 6.45±.27; N=20), *msi^1/1^* (green; 8.04±.23; N=28), transheterozygote *msi^1/2^* flies (pink; 8.73±.27; N=22), C587GAL4>+ (black; 6.11±.26) and C587GAL4>msi^RNAi^ (7.2±.25). Welch’s two-tailed t-tests reveal a significant increase in the average number of cysts in both *msi* null and *msi^1/2^* mutants compared to control *msi^1/+^* germaria (p<0.0001 in both cases) and in C587-GAL4>+ control germaria compared to germaria dissected from C587-GAL4>msi^RNAi^ flies (p=.0012). (D-E) High magnification single-plane confocal micrographs show colliding cysts (pink arrow, dotted outline) in ovaries dissected from *msi^1/1^* and *msi^1/M1^* mutants. (F-F’) Low magnification confocal micrographs of ovaries dissected from *msi^1/M1^* and *tj*-GAL4>msi^RNAi^ flies. Pink arrow points to fused egg chambers with two oocytes labelled with C(3)G. Scale bars, 20μm.

Since Msi is expressed in both somatic and germ cells, we next sought to determine whether swollen *msi* mutant germarial cysts resulted from somatic or germ cell abrogation of Msi function. To this end, *C587-GAL4*, a driver known to drive expression in EC and FSCs (42), coupled with an *UAS-msi^RNAi^* transgene (Supplementary Fig. 1B-C) was used to knockdown Msi function in somatic cells. Morphological analysis by immunofluorescence using anti-Vasa and anti-1B1 antibodies was then performed on ovaries dissected from 2-3 day old *C587-GAL4>msi^RNAi^* and *C587-GAL4/+* adults. Our results revealed a significant increase in the number germarial cysts in *C587-GAL4>msi^RNAi^* ovaries when compared to control *C587-GAL4* ovaries (Fig. 2B-C) suggesting a functional requirement for Msi in somatic cells of the ovary to regulate early ovary morphogenesis.

Along with the discovery of excess germline cysts upon loss of Msi function, immunofluorescence analysis of whole *msi^1/1^* mutants revealed other morphological defects. In about 32% of the *msi^1/1^* ovaries analysed (N=28), germline cysts in region 2a-2b appeared to exhibit a cyst collision phenotype (Fig. 2D). Cyst collision has been described as a process that can occur when forward moving germline cysts, or backward sliding cysts collide into the adjacent cyst (43). Cyst collision can transpire from the modification of the equilibrium between germline and somatic forces in germline cyst progression. Decreasing germline contractility or adhesion, or blocking somatic cell movement, can lead to germline cyst collision. Both scenarios have been shown to lead to the development of compound egg chambers (43). Since Msi function is abrogated in both the germline and somatic cells of *msi^1/1^* mutants, we assayed *msi^1/M1^* transheterozygote mutants for evidence of cyst collision. Msi function in *msi^1/M1^* flies would only be perturbed in somatic cells and the heteroallelic combination would circumvent any possibility of phenotypic effects caused from a second site mutation. Immunofluorescence analysis of *msi^1/M1^* germaria labelled with Vasa in combination with Fas 3 revealed evidence of cyst collision in 30% (N=60) of *msi^1/M1^* germaria analysed (Fig. 2E). Since colliding cysts have been shown to result in the formation of compound egg chambers (43), we labelled *msi^1/M1^* ovaries with the synaptonemal complex marker C(3)G (44), which revealed the presence of compound egg chambers containing 2 oocytes in 10% of egg chambers analysed (N=40) (Fig. 2F). We confirmed that the presence of compound egg chambers was due to somatic loss of Msi function by driving *UAS-msi^RNAi^* expression from the somatic cell driver *tj-GAL4*. In ovaries dissected from *tj-GAL4>UAS-msi^RNAi^* adults, approximately 10% of egg chambers (N=42) were compound chambers with 2 oocytes (Fig. 2F’).

Msi is expressed in all somatic cells in early oogenesis and in the germline. However, our observations of excess germ cells and germline cysts, along with cyst collisions, appeared confined to region 2a/b of *msi* mutant ovaries. We confirmed the number of GSCs and cystoblasts expressing phospho-Mad (the expression of which is restricted to these cell types (42) was not significantly different between control and *msi^1/1^* mutant ovaries (Supplementary Fig 4A-A’). Excess cysts were observed in regions 2a/b and 3, posterior to the Bag-of-Marbles (Bam) antibody expression domain, which labels 2-4 cell germline cysts (45) (Supplementary Fig. 4B-C). Taken together, our results demonstrate a functional requirement for Msi in region 2-3 somatic cells in the ovary in controlling early germline cyst morphogenesis.

Morphological analysis of *msi* mutants has uncovered a function for Msi in region 2-3 somatic cells of the germarium but we have not uncovered any evidence to support a requirement for the shorter isoforms of Msi in female germ cell maintenance or differentiation. Given the high level of Msi antibody expression observed in the GSCs of the adult ovary (Fig. 1B-C), we expected clonal analysis to reveal a functional requirement for Msi in the maintenance of GSCs comparable with what we previously observed in the testis (18). To further investigate this, we generated Flp-mediated FRT (Flp-FRT) *msi* loss of function GSC clones marked by the absence of GFP utilising two different *msi* mutant alleles, *msi^1^* and *msi^2^*, and measured whether the observed frequency of *msi* mutant GSC clones generated could be maintained over time. At 7 days post clone induction (PCI), the percentage of germaria containing at least one control GSC clone (56.7%, N=97) was not significantly different to the percentage of germaria containing at least one *msi^1^* (54.16%, N=96) or *msi^2^* (55.67%, N=97) mutant GSC clone at the same time-point (Fig. 3A). At 21 days PCI, no significant reduction in the percentage of germaria containing control (54.16%, N=97), *msi^1^* (50.98%, N=102) or *msi^2^* (53.00%, N=100) GSC clones was observed (Fig. 3A). These findings surprisingly revealed that, unlike the testis, there is no intrinsic requirement for the shorter Msi isoform/s in the maintenance of GSC identity in the adult *Drosophila* ovary.

**Figure 3.**
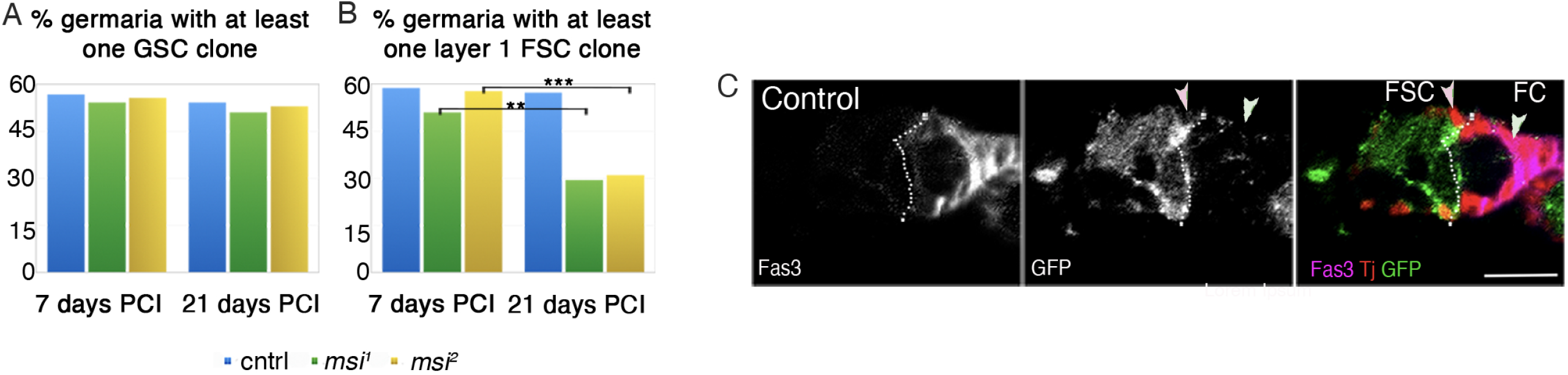
Msi is required for FSC maintenance, but not GSC maintenance. (A) The percentage of germaria containing at least on negatively marked control (blue), *msi^1^* (green) or *msi^2^* (yellow) GSC clone generated by Flp-FRT mediated recombination at 7 days PCI (56.7%, N=97; 54.16%, N=96 and 55.67%, N=97 respectively) and 21 days PCI (54.16%, N=96; 50.98%, N=102 and 53%, N=100 respectively). No significant differences were observed between genotypes as measured by Fisher’s exact test. (B) The percentage of germaria containing at least on negatively marked control (blue), *msi^1^* (green) or *msi^2^* (yellow) layer 1 FSC clone generated by Flp-FRT mediated recombination 7 and 21 days PCI. A significant reduction in the frequency of *msi^1^* (29.41%, N=102; Fisher’s exact test p=.0023) and *msi^2^* (31%, N=100; Fisher’s exact test, p=.0002) mutant FSC clones at 21 days PCI was observed. (C) Single-plane confocal micrographs showing Traffic Jam (Tj, red) positive, GFP-negative clones generated by Flp-FRT. A FSC clone (pink arrowhead) and FC clone (pale green arrowhead)) are depicted. Scale bars, 10μm. Dotted line represents the Fas3 expression boundary.

Since we discovered a functional requirement for Msi in region 2-3 somatic cells of the germarium to regulate cyst morphogenesis, we sought to determine whether Msi may play a role in the regulation or maintenance of FSCs, which also reside in this region. The percentage of germaria containing at least one Tj-positive but GFP-negative *msi^1^, msi^2^* or control layer 1 FSC clone at the 2a/b boundary was compared at 7- and 21-days PCI (Fig. 3C). After 7 days PCI, at least one layer 1 control FSC clone was observed in 58.76% (N=97) of germaria, not significantly different to the percentage of germaria containing at least one *msi^1^* (51.04%, N=96) or *msi^2^* mutant FSC clone (57.73%, N=97) at the same time-point (Fig. 3B). By 21 days PCI, the percentage of germaria containing at least one control FSC clone remained relatively unchanged (57.29%, N=96). In contrast, we observed a significant reduction in the percentage of germaria containing a at least one layer 1 *msi^1^* (29.41%, N=102; Fisher’s exact test p=.0023) or *msi^2^* (31.00%, N=100; Fisher’s exact test p=.0002) mutant FSC clones at 21 days PCI compared to 7 days (Fig. 3B), revealing a requirement for Msi in layer 1 FSC maintenance. Together, our results demonstrate an isoform specific requirement for Msi in germarial somatic cells to support cyst morphogenesis and maintenance of FSC identity.

Morphological analysis of whole *msi* mutants revealed a cyst collision phenotype in region 2a-2b of the germaria in approximately one third of ovaries examined (Fig. 2D-E). In this region, somatic cells directly anterior and adjacent to layer 1 FSCs have been described as proliferatively active layer 2-3 FSCs (16). Also in this region reside posterior escort cells, otherwise known as inner germarial sheath (IGS) cells (11, 12). In our clonal analysis, we did observe the presence of negatively marked region 2a-2b control somatic cell clones in cells that corresponded to FSC layers 2 and 3 at both 7- and 21- days PCI (Supplementary Fig. 5A-B). While the percentage of germaria containing somatic control clones in this region remained similar at both 7- and 21- days PCI, we observed a significant difference in the percentage of germaria containing at least one layer 2 or 3 *msi^1^* (Fisher’s exact test p=.03) or *msi^2^* (Fisher’s exact test p=.01) clone between 7- and 21- days after PCI (Supplementary Fig. 5B). Moreover, some *msi* mutant clones appeared to be aberrantly positioned on the outer edge of the ovary (Supplementary Fig. 5C). Loss of *msi* mutant somatic cell clones from region 2a/b led to the question of whether mutant clones were dying. To test this, we generated GFP-labelled *msi^1^* mutant FSC clones by MARCM (Mosaic Analysis with a Repressible Cell Marker) to visualise whether GFP marked clones co-expressed the cell death marker Dcp-1 (*Drosophila* caspase-1). Of the 27 *msi^1^* mutant layer 1-3 FSC clones analysed 12 days PCI, none expressed Dcp-1, suggesting that *msi* mutant clones were not being lost due to cell death (Fig. 4A). However the morphology of CD8::GFP labelled *msi^1^* FSC clones appeared to differ from wild-type counterparts. FSCs in layers 1-3 have been shown to extend processes, or axon-like projections, across the entire germarium (16, 46). Of the 27 germaria carrying a GFP marked *msi^1^* mutant FSC clone, 11 displayed aberrant, shortened projections (Fig. 4B-C). In comparison, we only observed 3 control GFP clones with aberrant extensions. Shortened processes in somatic cells in region 2a is a hallmark of escort cell morphology (16) and it is possible that loss of Msi function could be causing some FSCs to adopt an escort cell fate. This would be consistent with the known intrinsic requirement of Msi to maintain stem cell self-renewal capabilities in *Drosophila* GSCs (18). Unfortunately, we have been unable to test this hypothesis owing to a lack of identification of FSC-specific drivers and markers to interrogate this question.

**Figure 4.**
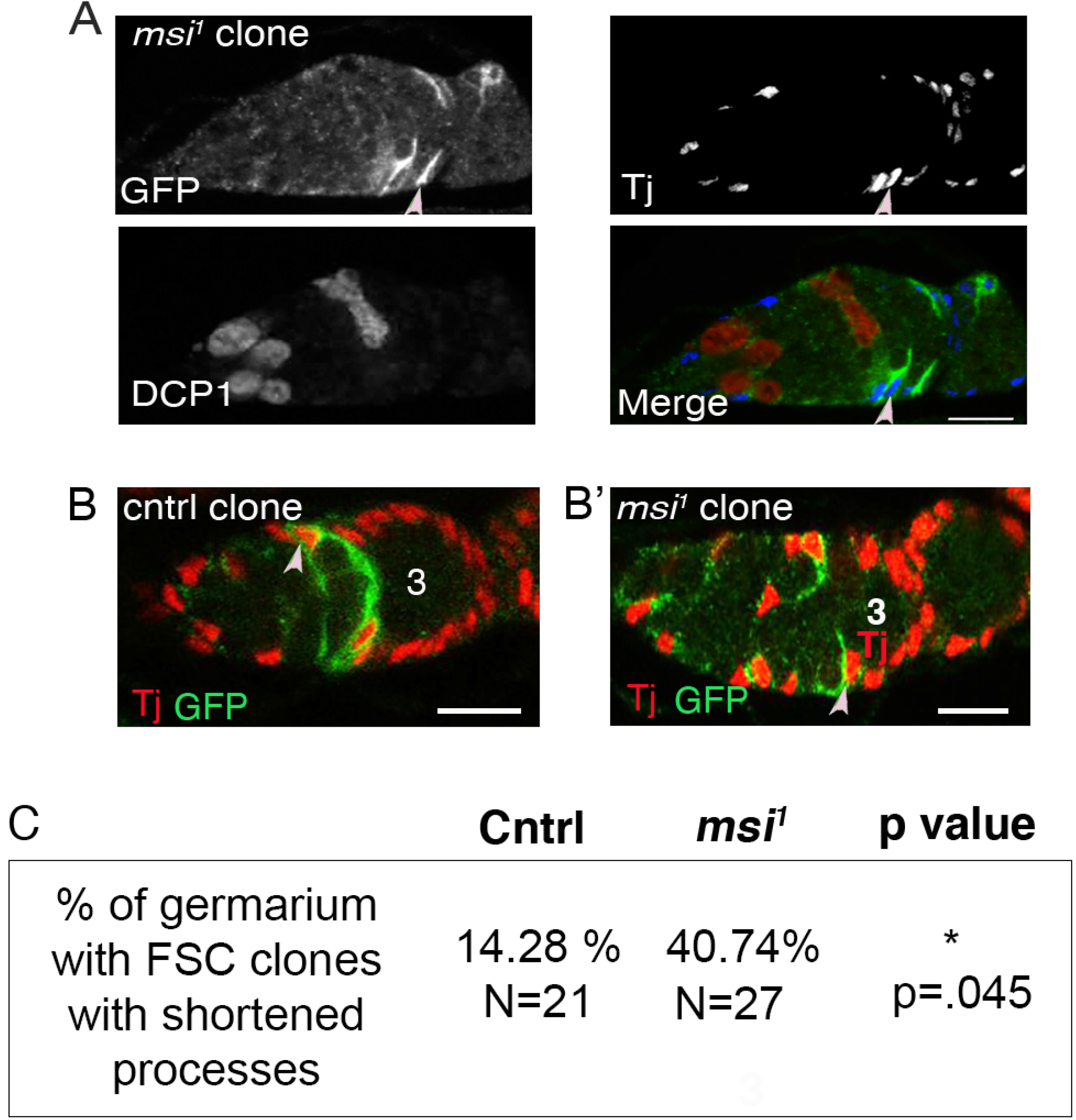
*msi* mutant FSC clones do not express Dcp-1 but display aberrant morphology. (A) Single-plane confocal micrographs showing Tj-positive (blue in merged panel) and GFP-positive MARCM *msi* null FSC clone (green in last panel). This image depicts Dcp-1 positive germ cells dying as a control for Dcp-1 antibody labelling. (B) Single-plane confocal micrograph showing a control, layer 1 GFP-positive FSC clone (light pink arrowhead) with an axon-like projection extending across the germarium. (B’) Single-plane confocal micrograph showing a *msi* null, layer 1 GFP-positive FSC clone (light pink arrowhead) with a shortened, aberrant projection. (C) Table represents the % of control and *msi^1^* mutant 10 day old FSC MARCM clones with shortened axon-like processes. p-value was calculated in Prism 9 for MacOS using Fisher’s exact test. Scale bars, 10μm. 3 represents a region 3/stage 1 egg chamber.

The aberrant morphology of *msi* mutant FSCs concomitant with loss of somatic cells from region 2a/b is suggestive of cell differentiation defects, perhaps at the expense of proliferative capacity. Because FSCs in layers 1-3 in region 2a/b of the germarium have been defined as proliferatively active, we asked whether loss of Msi function would lead to cell cycle defects. To this end, we used the *Drosophila* Fluorescence Ubiquitin-based Cell Cycle Indicator (Fly-FUCCI) system in which degradable versions of GFP::E2f1 and RFP::CycB fluorescently label cells in G1, S and G2 phases of interphase (47) to determine whether *msi* mutant somatic cells were cycling normally. In the fly FUCCI system, G1 is distinguished by GFP::E2F1 labelling in the absence of RFP::CycB. G2 cells appear yellow as they are labelled with both GFP and RFP, and cells in S phase are labelled by RFP alone. *109-30-GAL4* was used to drive a GAL4 responsive UAS-fly-FUCCI transgene (UAS-FUCCI) in somatic cells encompassing FSC layers 1-3 and escort cells abutting the posterior of region 2a of the germaria (11) (Fig. 5A-B). Counting somatic cells within the boundary of region 2a and region 2b and excluding mature follicle cells, knockdown of Msi by RNAi resulted in a significant reduction in the average number of somatic cells in G1 phase per germaria (Fig. 5C). Chi-square analysis also revealed a reduction in the proportion of *109-30-Gal4>UAS-msi^RNAi^* mutant germaria containing at least one cell in G1 phase (p=.005; Fig. 5C’). Knockdown of Msi by RNAi also resulted in a significant increase in the average number of cells in G2 per germaria when comparing to controls (t-test p<.0001; Fig. 5D) and a significant increase in the total number of germaria exhibiting at least one somatic cell in G2 phase (Chi-square p<.0001; Fig. 5D’). Together our results suggest that loss of Msi function from somatic cells within region 2a-2b of the germarium leads to a lag in G2 phase of the cell cycle and supports the overall finding that Msi is required to maintain the identity of somatic stem cells and also to support the function of escort cells in this region.

**Figure 5.**
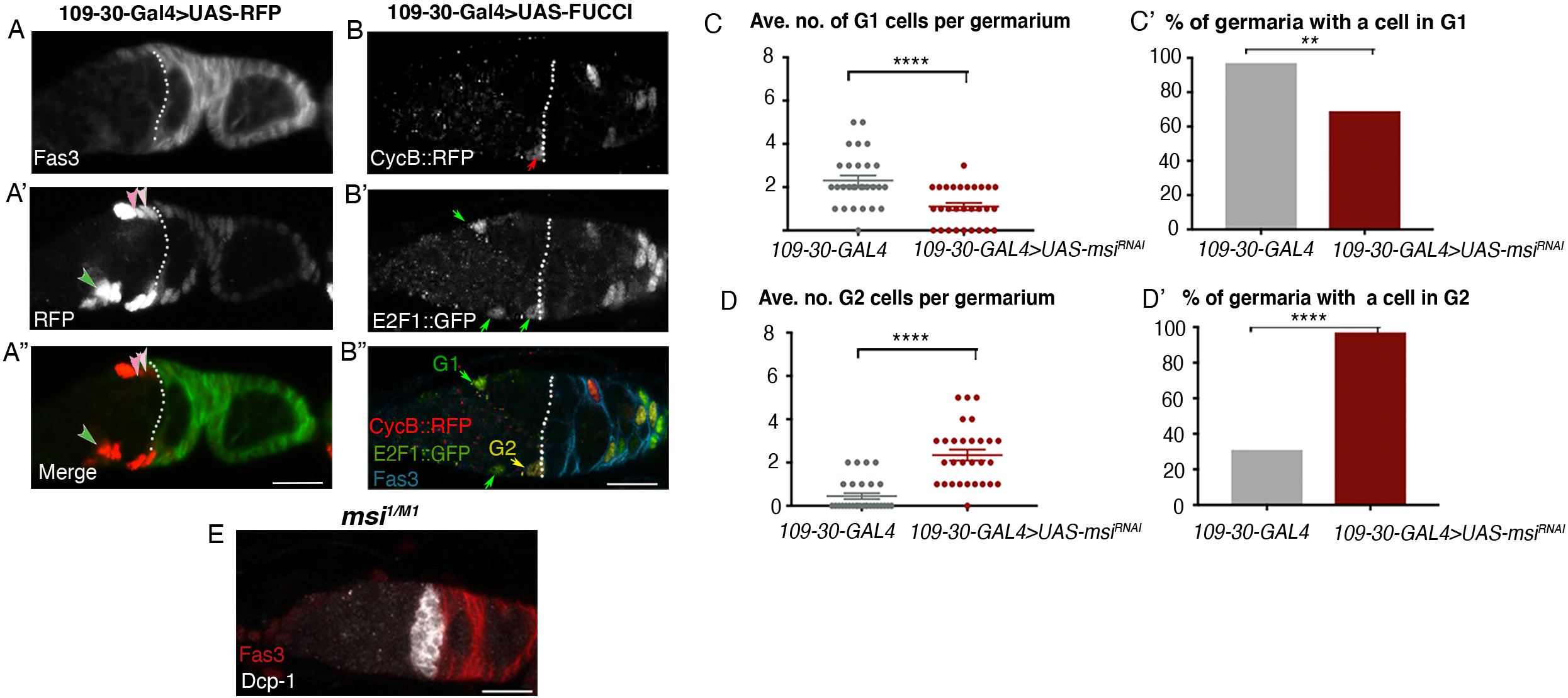
Loss of Msi function in somatic cells causes cell cycle defects and an increase in dying germline cysts in region 2a/b of germaria. (A-A”) Confocal micrograph (projection of 3 planes from a z stack) showing a representative image of a *109-30-Gal4>UAS-RFP* adult germarium. Dotted line represents the Fas3 expression boundary. A FSC in each of layers 1 and 2 (light pink and medium pink arrowheads respectively) and a posterior EC (green arrowhead) that are expressing RFP are labelled in A’ and A”. (B) Confocal micrograph showing a representative image of a *109-30-Gal4>UAS^FUCCI^* adult germarium. A CycB::RFP-positive cell (red arrow), E2F1::GFP positive cells (green arrows), A G2 cell (yellow arrow in B”) and G1 cells (green arrow in B”) anterior to the Fas3 expression boundary (dotted line) are labelled. (C) Scatterplot showing a significant difference in the average number of cells (± SEM) in G1 within region 2a −2b of germaria dissected from *109-30-Gal4* (2.31 ± 0.23, N=29) and *109-30-Gal4>UAS-msi^RNAi^* (1.18 ± 0.16, N=29)(t-test p<.0001) adults. (C’) Column graph showing a significant decrease in the % of *109-30-Gal4>UAS-msi^RNAi^* germaria with at least one cell in G1 (69%; N=29; Chi-square p=.005) when compared to *109-30-Gal4* germaria (97%). (D) Scatterplot showing a significant difference in the average number of cells (± SEM) in G2 within region 2a-2b of germaria dissected from *109-30-Gal4* (0.45 ± 0.14, N=29) and *109-30-Gal4>UAS-msi^RNAi^* (2.35± 0.25, N=29)(t-test p<.0001) adults. (D’) column graph showing significant increase in the % of *msi^RNAi^; 10930^Gal4^* (97%; N=29; Chisquare p<.0001) mutant germaria with the presence of at least one cell in G2 compared to control germaria (31%). (E) Single-plane confocal micrograph of *msi^1/M1^* ovary labelled with Dcp-1 (white) and Fas3 (red). Scale bars, 10μm.

Our findings have established an isoform specific function for Msi in maintaining FSC fate and supporting germline cyst morphogenesis in early oogenesis. Specifically, *msi* mutant somatic cells in region 2a/b display aberrant morphology and appear to lag in G2 phase of the cell cycle. While there is no evidence to suggest that *msi* mutant somatic cells die, clonal analysis demonstrated that *msi* mutant FSCs fail to be maintained in this region of the germarium. Furthermore, an accumulation of germline cysts in region 2-3 of the germarium, cyst collisions and the formation of compound egg chambers suggest that somatic cells fail to properly interact with, and support, the developing germline. Further evidence to support this comes in the way of analysis of cell death in whole *msi* mutants. When analysing cell death using an antibody to detect Dcp-1, we observed roughly a twofold increase in the number of germline cysts dying in a *msi^1/M1^* (11/31) whole mutant germarium compared to a *w^1118^* control germarium (5/32) (Fig. 5E). Normally, cell death in region 2b of the germarium occurs sporadically in well fed flies (reviewed in (48)). In *msi^1/M1^* mutants, 9 of the 11 dying cysts in *msi^1/M1^* mutants were in region 2a/b, supporting the hypothesis that *msi* mutant somatic cells in this region are defective in providing the necessary signals to the germline to fully support germline cyst progression in early oogenesis.

The formation and separation of egg chambers in early oogenesis relies upon correct differentiation of FSCs and their progeny within the germarium. Egg chambers are separated by 5-8 flattened disc-like somatic stalk cells which have differentiated from the pool of follicular precursor cells and connect the anterior end of a more mature egg chamber with the posterior end of a younger egg chamber (15). Stalk cells, along with disc-like terminal filament cells and cap cells, express high levels of Lamin C (Lam C) (Fig. 6A). We asked whether stalk cell formation occurred normally in *msi^1/M1^* mutant ovaries by labelling whole ovaries with an antibody to detect Lamin C expression. While Lamin C-positive stalk cells were observed in *msi^1/M1^* mutant ovaries (Fig. 6B) and the number of stalk cells did not appear to be affected by loss of Msi protein function, we surprisingly observed an up-regulation of Lamin C expression in FSCs, FCs and some ECs in the germarium (Fig. 6B-B’). Up-regulation was consistently observed in region 2a/b of the germaria (Fig. 6B), with occasional region 1 escort cells displaying increased Lamin C levels (Fig. 6B’). These results support the idea that loss of Msi function from somatic cells within this region results in altered differentiation potential of FSCs within the germarium and that Msi is required to maintain somatic stem cell identity. Our results also point to a functional requirement for Msi in a specific subpopulation of ECs in region 2a/b of the germarium to support germline cyst morphogenesis.

**Figure 6.**
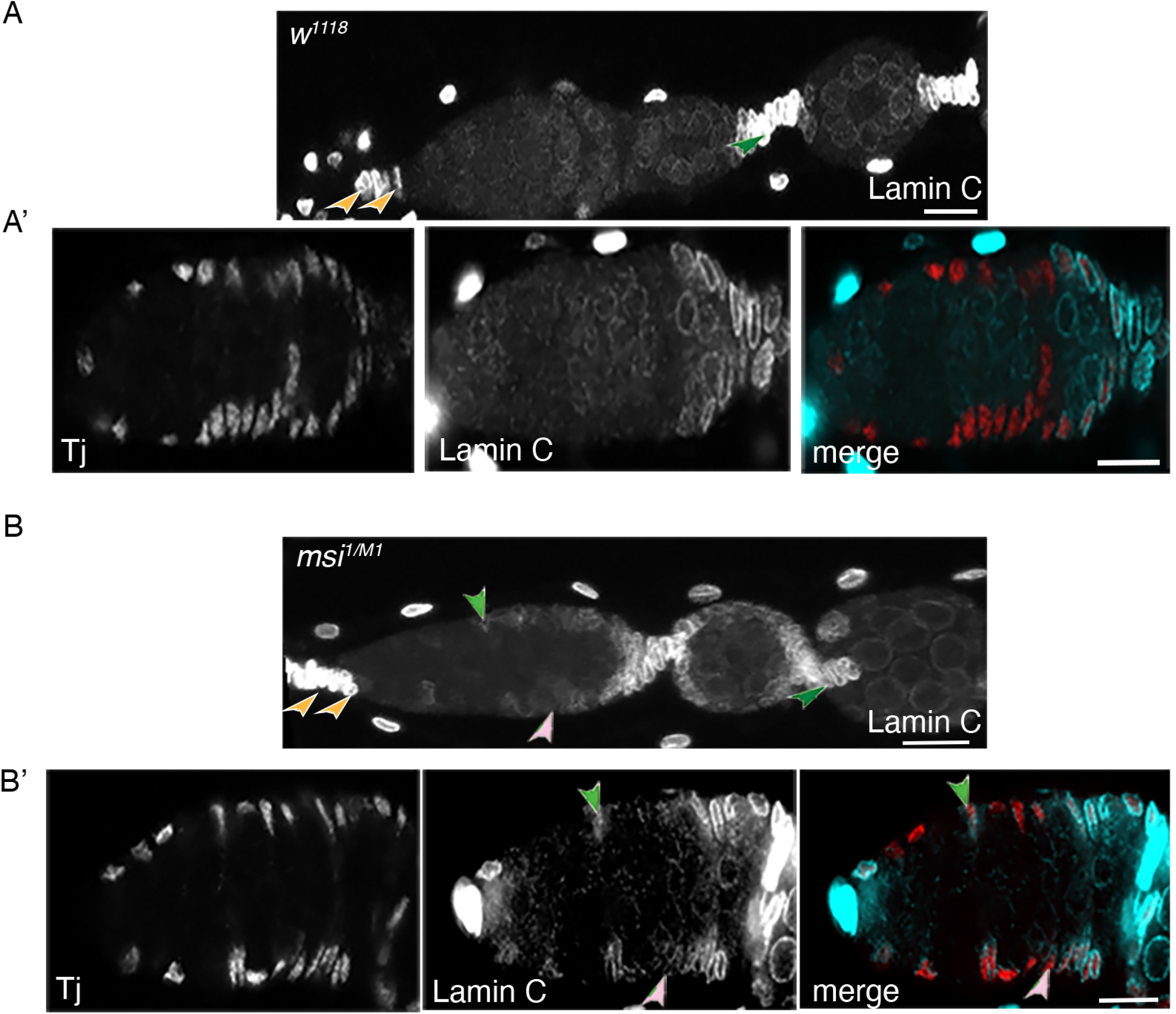
Losing Msi function from somatic cells results in mis-expression of Lamin C. (A-A’) Confocal micrographs (projection of 3 planes from a z stack) showing normal Lamin C expression in TFs and CCs (orange arrowheads) and stalk cells (green arrowhead) in a wild-type germarium. (B-B’) Confocal micrographs (projection of 3 planes from a z stack) showing up-regulation of Lamin C in somatic cells in germaria dissected from *msi^1/M1^* adults. A layer 1 FSC (light pink arrowhead) and posterior EC (green arrowhead) expressing Lamin C are labelled. Scale bars, 10μm.

## Discussion

*Drosophila* Msi was first identified for its role in determining the fate outcome of ectodermal sensory organ precursor cells (41). Subsequently, in flies, Msi has been shown to play roles in photoreceptor cell differentiation (36) and in crystal cell determination (49). In *Drosophila* stem cell populations, Msi has been shown to be required for the maintenance of spermatogonial germline stem cells (18) and can modulate intestinal stem cell proliferation in guts damaged from exposure to radiation (22). These studies highlight the importance of *Drosophila* Msi in the context of cell fate determination and stem cell maintenance in many cell types.

*Drosophila msi* encodes multiple polypeptides (Fig. 1G) (35). To date no study has probed whether these isoforms play specific or different roles in development. Our analysis has revealed that Msi isoforms exhibit divergent patterns of expression in the germline and soma of both the adult *Drosophila* testis and ovary. In both cases the longer protein isoforms were only detected in the somatic cells. Surprisingly we found that there is no functional requirement for shorter Msi isoform/s Msi-RA and/or Msi-RH in GSC maintenance in the female despite this/these isoforms being highly expressed in female GSCs and intrinsically required to maintain GSC identity in the *Drosophila* testis (18). Since Msi is an RBP and its target mRNAs remain unknown, this disparity could simply reflect differences in the genetic machinery required to regulate GSC identity in males and females. Sex specific differences in RBP function have certainly been described with other RBPs such as Pumilio (Pum), Nanos (Nos) and Held Out Wings (How). Pum functions with Nos in female GSCs to prevent GSC differentiation into a daughter cystoblast (50, 51). Despite spermatogonial stem cells expressing Nos, there is no evidence that Pum is expressed in GSCs of the testis and no functional role for Pum has been discovered for male GSC maintenance. How is expressed and required in both male and female GSCs, but the mechanism of action differs (52, 53). In males, How was shown to be required to maintain Cyclin B (CycB) levels in GSCs (52). Loss of *how* resulted in decreased levels of CycB, leading to a delay in G2 phase of the cell cycle. Consequently, GSCs were unable to enter mitosis and were lost from the niche via cell death. In contrast, female GSCs were dependent upon How function due to its requirement to repress Bam expression (53), since loss of *how* led to *bam* deregulation in GSCs and subsequent premature differentiation of these cells. In female GSCs, no evidence for the regulation of CycB levels by How was found, with this result reflecting differential use of the How RBP in male and female GSCs.

Despite not finding a requirement for short Msi isoforms (Msi-RA and/or Msi-RH) in maintaining GSC function in the ovary, we did demonstrate that one (or more) of the longer Msi isoforms (Msi-RB, Msi-RC, Msi-RD, Msi-RE or Msi-RF) is required to regulate the differentiation outcome of germarial FSCs. We showed that *msi* null and hypomorphic mutant FSC clones, generated by Flp-mediated recombination, were not maintained in the germarium since FSCs in layer 1 and the adjacent layers 2 and 3 were lost over time. MiMIC analysis and expression data showed that only the longer Msi isoforms are expressed in somatic cells of the ovary, therefore demonstrating a divergence of Msi isoform function in a context-dependent manner for the regulation of different stem cell populations. Our finding that Msi is required to maintain the epithelial stem cell fate is consistent with the known function of its vertebrate orthologues, Msi-1 and Msi-2. Vertebrate Msi-2 has been shown to play a critical role in hematopoietic cell fate and lineage bias (29). Vertebrate Msi proteins have been shown to be associated with epithelial cell state in several cancer types, most notably breast cancer, with loss of Msi leading to loss of epithelial cell identity (38). In the intestinal epithelium, a requirement for vertebrate Msi proteins in maintaining quiescent intestinal stem cells has been demonstrated (54). The discovery that Msi is required for the regulation of an epithelial stem cell population in *Drosophila* provides a research model to investigate the molecular mechanisms that underpin the maintenance of epithelial stem cells in a context dependent manner.

An interesting aspect arising from our research was the discovery that Msi is required in a distinct population of somatic ECs to support early germline cyst progression in oogenesis. Several studies have revealed morphological and functional differences in ECs depending on their position within the germarium (7). Recent single cell sequencing analyses, in attempts to provide molecular atlases of the *Drosophila* ovary, have demonstrated the existence of at least two EC subgroups (11, 13), with one study claiming that there is as many as four distinct EC subpopulations (12). These analyses have uncovered functional differences between EC populations, with anterior ECs acting on GSCs and cystoblasts to support synchronous cell division, while more posterior ECs regulate soma-germline cell adhesion and the transition from 16-cell cyst-to-egg chamber formation. In our study we found no evidence to suggest that Msi function is required in anterior ECs. In *msi^1/1^* mutant ovaries, early germline development progressed normally. We observed that the number of GSCs and cystoblasts expressing phospho-Mad was not significantly different between control and *msi^1/1^* mutant ovaries. Furthermore, the excess cysts that were observed in *msi* mutants was restricted to regions 2a/b and 3, posterior to the Bag-of-Marbles (Bam) antibody expression domain, which labels 2-4 cell germline cysts. Shi and colleagues (2021) observed a similar swollen germarial phenotype to that which we observed in whole *msi* mutant ovaries, including the accumulation of 8 cell cysts in germaria upon ablation of posterior ECs. In context of this recent literature, our results indicate a specific requirement for Msi function in a subpopulation of ECs and likely reflects the requirement for Msi to bind target mRNAs which have expression limited to somatic cells within this domain of the germarium.

In our study we have identified a functional requirement for Msi in FSCs and posterior ECs in the adult *Drosophila* ovary. The demarcation between ECs, FSCs and FSC progeny is not well characterised and remains controversial. Several single-cell sequencing projects have failed to identify a distinguishable FSC population (11, 13). Shi and colleagues also noted that mitotic activity between FSCs and ECs was not sufficiently different, and these cells could not be separated using a methodology designed to identify stem cell populations. Others have demonstrated that posterior ECs can convert to FSCs, at least under starvation conditions (Rust et al., 2020). Further studies have demonstrated that FSCs can give rise to posterior ECs and even anterior ECs over time (11, 16). Given the mounting evidence that FSCs and posterior ECs are similar in their transcriptional profile and overlap in function, it is likely that the binding targets of Msi are the same in both cell types and Msi is required to maintain the fate and functionality of both FSCs and posterior ECs. Consequently, loss of Msi function from these cells results in their inability to support germline cyst progression and fate interchangeability. Indeed, loss of Msi function from somatic cells in the 109-30-Gal4 expression domain, which encompasses both FSCs and posterior ECs, results in a lag in the G2 phase of the cell cycle and an up-regulation of a differentiation marker Lam C. These cells are not lost from the germarium due to cell death, but mis-expression of Lamin C implicates a change in cell fate due to loss of Msi function. Since Msi is a translational repressor, we considered that *Lam C* mRNA may be a direct target of Msi. However, no Msi binding sequences are present in the 3’ UTR of *Lam C*, making it an unlikely target.

Our study has identified isoform specific requirements for Msi in stem cell maintenance in *Drosophila*. Mammalian Msi-2 is the closest Msi orthologue to *Drosophila* Msi and has four isoforms, all of which contain two RNA-recognition motifs (RRMs) required for binding mRNAs but differ in the N-terminus or C-terminus (34). Intriguingly, this is comparable to *Drosophila* Msi isoforms, where the RRMs are conserved between the isoforms but proteins differ at the N-terminus (Supplementary Fig. 2). Only recently, studies attempting to understand mammalian Msi-2 isoform function in development have begun to highlight the different roles these isoforms may play in tissue homeostasis. One study has shown that a truncated Msi-2 isoform lacks regulatory phosphorylation sites and is overexpressed in multiple cancers (55). Another more recent study has highlighted the differential expression patterns of Msi-2 isoforms in triple-negative breast cancer (TNBC) tissue and has demonstrated that downregulation of a predominant functional isoform (Msi-2a) is associated with TNBC progression (34). Vertebrate Msi proteins have been associated with regulation of mRNA stability, splicing and either enhancement or suppression of translation (38, 56–60). Phosphorylation of C-terminal serine residues has been demonstrated to regulate the switch between translational repression and activation, and this in turn can be regulated by Msi protein isoform usage (55). Future studies into the role of *Drosophila* Msi isoforms in development will add insight into how specific isoforms can differentially regulate stem cell behaviour in a sex- and cell-specific manner.

## Materials and Methods

### Fly Strains

All flies were raised on standard molasses-based food at 25°C except for Gal4 crosses, which were all carried out at 29°C. Fly stocks used in this study obtained from the Bloomington stock centre (Indiana) include *w^1118^, msi^1^, msi^2^*, Mi{MIC}msi^M101988^ (*msi^M1^;* BL33097), Mi{PT-GFSTF.2}msi^M100977-GFSTF.2^ (Msi-GFP; BL61750), 109-30-Gal4, UAS-mcD8::GFP (CD8::GFP) reporter on 1^st^ and 3^rd^ chromosomes, UAS-FUCCI transgenes (BL55110, BL55111), *FRT82BUbi-GFP*, Frt42D;FRT82B and *hh*-lacZ. *frt82Bmsi^1^* and *frt82Bmsi^2^* were previously made in our laboratory (18). MARCM82B flies (hsflp, UAS-GFP::CD8;; tubulin-GAL4, FRT82B tubP-GAL80) were a gift from the Quinn lab (Australia National University). The X chromosome *msi^RNAi^* strain was obtained from the Vienna Drosophila Resource Center (VDRC #11784). *tj*-GAL4 (DGRC104055) was obtained from Kyoto Stock Center (Japan). c587-GAL4 and *Fax-GFP* lines were gifts from the Xie lab (Stowers Institute for Medical Research, Missouri, USA).

### Immunostaining and image analysis

Appropriately aged females were dissected in PBS, fixed for 20 minutes in 4% Formaldehyde in PBT (PBS +.2% Triton X-100 (Sigma)), washed for 3x 10 minutes in PBT and then incubated for 45 minutes in PBTH (5% Normal Horse Serum diluted with PBT). Ovaries were then incubated overnight at 4°C in primary antibodies diluted in PBT. Samples were washed a further 3×10 minutes in PBS before secondary antibody incubation was carried out for 2 hours at room temperature in PBT. Samples were washed for a further 3×10 minutes before ovaries were mounted on slides in Prolong™ Gold Antifade Reagent with Dapi (Invitrogen). Antibodies used in this study include 1:10 rat anti-Msi (gift of H. Okano, Keio University), 1:20 mouse anti-Fas3 (Developmental Studies Hybridoma Bank (DSHB)), 1:40 mouse anti-Lamin C (DSHB), 1:100 rabbit anti-DCP1 (Cell Signaling technologies), 1:100 goat anti-Vasa (Santa Cruz Biotechnology), 1:2000 chicken anti-GFP (Abcam), 1:500 rabbit anti-RFP (Invitrogen), 1:10 mouse anti-1B1 (DSHB), 1:1000 chicken anti-*β* galactosidase (Abcam), 1:10,000 guinea pig anti-Traffic Jam (gift of Dorothea Godt, University of Toronto) and 1:500 mouse anti-C(3)G (gift of Scott Hawley, Stowers Institute of Medical Research, Kansas City). Secondary antibodies (Donkey anti-Mouse Alexa Fluor 488/ 594/ 647, Donkey anti-Rabbit Alexa Fluor 594, Donkey anti-Rat Alexa Fluor 488/594) were obtained from (Thermo Fisher Scientific) and used at a dilution of 1:500. Donkey anti-guinea pig 594, and donkey-anti chicken 488 were obtained from Jackson Immuno Research Labs and used at a dilution of 1:500. Images were acquired on Zeiss LSM800 or LSM880 confocal microscopes as serial optical sections (z-stacks) optimized to acquire overlapping sections. FIJI/ ImageJ was then used to process images and add scale bars. Adobe photoshop 2021 was used to compile figure panels.

### Mosaic analysis

Negatively marked GSC and FSC clones were induced by Flp-mediated recombination at FRT sites in 2-3 day old females of genotypes *hs*-Flp/+; FRT82B *msi^1^*/FRT82B *Ubi*-GFP, *hs*-Flp/+; FRT82B *msi^2^*/FRT82B *Ubi*-GFP, *hs*-Flp/+; FRT42D/+;FRT82B/FRT82B *Ubi*-GFP. Well-fed females were heat shocked in a water bath at 37°C for 1 hour, twice a day for 2 consecutive days. Daily heat shocks were conducted 8 hours apart to aid recovery between heat shocks. Ovaries were dissected from 7 and 21 day old females (post heat shock) and stained with anti-GFP, anti-Traffic Jam and anti-Fas3. All GFP-negative but TJ positive clones at the 2a/2b boundary were counted as layer 1 FSC clones. All GFP-negative but TJ positive clones in the layers directly anterior and adjacent to layer 1 clones were counted as layer 2-3 clones. All GFP-negative but TJ positive clones anterior to layer 2-3 clones were counted as escort cell clones.

### MARCM analysis

GFP positive clones were generated in 2-3 old adult females of genotypes hsflp, UAS-GFP::CD8/+; tubulin-GAL4, FRT82B tubP-GAL80/FRT82B and hsflp, UAS-GFP::CD8 /+; tubulin-GAL4, FRT82B tubP-GAL80/FRT82B *msi^1^*. Females were subject to a 45 minute heat shock in a 37°C waterbath, twice a day, for two consecutive days, with sufficient time in between heat shocks to ensure recovery. Ovaries were dissected at 10 days post heat shock and stained with anti-GFP, anti-Traffic Jam and anti-DCP1, and imaged as previously described. FIJI was used to analyse the images and the Fisher’s exact test in Prism 9 for Mac OS was used to calculate p values.

### FUCCI analysis

Full genotypes of flies used in FUCCI analysis include 109-30-Gal4/UASp-GFP::E2f, UASp-mRFP1.NLS::CycB; *+/+* (control) and *msi^RNAi^/+;* 109-30-Gal4/+; UASp-GFP::E2f, UASp-mRFP1.NLS::CycB. Flies were raised at 25°C and shifted to 29°C upon eclosion for 3 days. Well-fed females were dissected and stained for detection of GFP and RFP. Serial overlapping optical sections were analysed in FIJI, with DAPI used in conjunction with the ROI manager to make sure to not duplicate cell counts. The minimal brightness threshold utilised for the RFP and GFP channels was 100.

### Statistics

Statistical analyses were performed using Prism 9 for Mac OS. p-value calculations for all statistical analyses are noted in Figure legends. All scatterplots are graphed showing the mean ± SEM.

## Supporting information

Supplemental data, methods and references

## Acknowledgments

The authors wish to thank Ting Xie, Leonie Quinn, Bloomington Drosophila Stock Center, Vienna Drosophila RNAi Center and the Australian Drosophila Biomedical Research Support Facility (OzDros) for provision of *Drosophila* strains. We also wish to thank Hideyuki Okano, Scott Hawley, Dorothea Godt and the Developmental Studies Hybridoma Bank for provision of antibodies. We are grateful for the assistance of the Biomedical Optical Microscopy Platform at the University of Melbourne. This work was conducted with the support of Australian Research Council Discovery Project Grants 120100224 to EM and GH and 170102379 to GH.

