## Supplemental data, methods and references for "MiMIC analysis reveals an isoform specific role for *Drosophila* Musashi in follicle stem cell maintenance and escort cell function"

### Supplemental Materials and Methods

#### Additional fly strains and antibodies

Additional fly strains used for the supplementary methods include *tub*-GAL4 (obtained from Bloomington Stock Center). Additional antibodies include Mouse anti-Bam 1:10 (DSHB) and rabbit anti-pMad (pSmad1/5, 41D10, 1:100; Cell Signalling).

#### pMad antibody staining protocol

Immunostaining was carried out as per the methods section of the paper except Sodium orthovanadate (1:100, Sigma) was used as a phosphatase inhibitor in the fixation step. For this protocol, fixation was undertaken on ice.

#### Testis antibody staining protocol.

3 day old adult testes were dissected and stained as per our ovary protocol.

#### Mosaic analysis

Negatively marked clones depicted in Supplementary Figure 5 were generated as per methods section in the paper. All GFP-negative but TJ positive clones at the 2a/2b boundary were counted as layer 1 FSC clones. All GFP-negative but TJ positive clones in the layers directly anterior and adjacent to layer 1 clones were counted as layer 2-3 clones. All GFP-negative but TJ positive clones anterior to layer 2-3 clones were counted as escort cell clones.

**Image analysis**

Images were acquired on Zeiss LSM800 or LSM880 confocal microscopes as serial optical sections (z-stacks) optimized to acquire overlapping sections. FIJI/ ImageJ was then used to process images and add scale bars. Fiji/ImageJ was also used to create the orthogonal view image in Supplementary Figure 5. The adult fly image was captured on a dissecting microscope (Olympus) with a DP20 camera attachment (Olympus). Image was processed using Adobe Photoshop.

**Sequence alignment**

Sequence alignment was conducted using Clustal O (1.2.4).

**Statistics**

Statistical analyses for Supplementary Figures were performed using Prism 9 for Mac OS. p-value calculations for all statistical analyses are noted in Figure legends. All scatterplots are graphed showing the mean ± SEM.

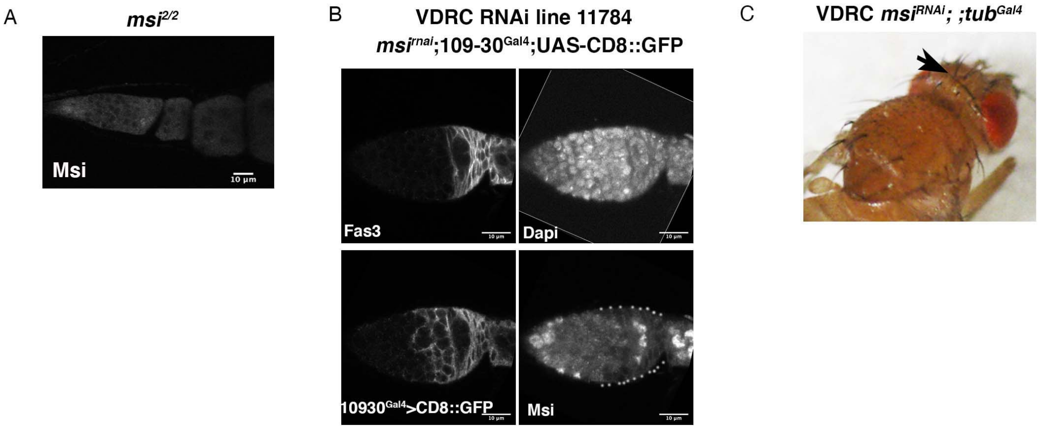


**Supplementary Figure 1. Musashi antibody specificity and RNAi knockdown efficiency.** (A) Confocal micrograph of Msi antibody staining on *msi*$^{2/2}$ germaria showing very little Msi expression. (B) Confocal micrographs of germaria where the UAS-*msi*$^{RNAi}$ transgene was driven from the *109-30-Gal4* driver. The expression domain of *109-30-Gal4* is marked by GFP (third panel). Knockdown of Msi antibody expression in the *109-30-Gal4* expression domain was observed (dotted outline of last panel). The white outline of the Dapi channel represents where the image was rotated to display the correct orientation. (C) Knockdown of the RNAi line from *tubulin*-Gal4 *(tub*$^{Gal4}$*)* results in the double bristle phenotype (arrow) originally identified by Nakamura and colleagues (1). Scale bars, 10µm.

.

CLUSTAL O(1.2.4) multiple sequence alignment

```
Msi_PE        MLFENPAVAAKLPFPYNVPPPLQAAAAAAAAVPNLR----------------SVSEMNAT 44
Msi_PD        MLFENPAVAAKLPFPYNVPPPLQAAAAAAAAVPNL------------------------- 35
Msi-PBCF      MLFENPAVAAKLPFPYNVPPPLQAAAAAAAAVPNLRFQTPIKAFACLTAATRSVSEMNAT 60
Msi-PA        ---------------------MHALQEG---ATVLHHQQPPPP----TSGEDHLLTADSF 32
Msi_PH        ------------------------------------------------------------ 0

Msi_PE        SLYAGNPMENAAAAAAAAAAGLIDPHHNRDLHQALVASIANNSVAAIGGGLTTAAVLKSA 104
Msi_PD        ----SNPMENAAAAAAAAAAGLIDPHHNRDLHQALVASIANNSVAAIGGGLTTAAVLKSA 91
Msi-PBCF      SLYAGNPMENAAAAAAAAAAGLIDPHHNRDLHQALVASIANNSVAAIGGGLTTAAVLKSA 120
Msi-PA        FYARSNPMENAAAAAAAAAAGLIDPHHNRDLHQALVASIANNSVAAIGGGLTTAAVLKSA 92
Msi_PH        -------MENAAAAAAAAAAGLIDPHHNRDLHQALVASIANNSVAAIGGGLTTAAVLKSA 53
                     *****************************************************

Msi_PE        AQQSQQAVQQNQNAVVVTPGLEQPKQEPAQQAALALLKENVNASAGAGQNNGQAAMGGSN 164
Msi_PD        AQQSQQAVQQNQNAVVVTPGLEQPKQEPAQQAALALLKENVNASAGAGQNNGQAAMGGSN 151
Msi-PBCF      AQQSQQAVQQNQNAVVVTPGLEQPKQEPAQQAALALLKENVNASAGAGQNNGQAAMGGSN 180
Msi-PA        AQQSQQAVQQNQNAVVVTPGLEQPKQEPAQQAALALLKENVNASAGAGQNNGQAAMGGSN 152
Msi_PH        AQQSQQAVQQNQNAVVVTPGLEQPKQEPAQQAALALLKENVNASAGAGQNNGQAAMGGSN 113
              ************************************************************

Msi_PE        KSGSSGRSTPSLSGGSGSDPAPGKLFVGGLSWQTSSDKLKEYFNMFGTVTDVLIMKDPVT 224
Msi_PD        KSGSSGRSTPSLSGGSGSDPAPGKLFVGGLSWQTSSDKLKEYFNMFGTVTDVLIMKDPVT 211
Msi-PBCF      KSGSSGRSTPSLSGGSGSDPAPGKLFVGGLSWQTSSDKLKEYFNMFGTVTDVLIMKDPVT 240
Msi-PA        KSGSSGRSTPSLSGGSGSDPAPGKLFVGGLSWQTSSDKLKEYFNMFGTVTDVLIMKDPVT 212
Msi_PH        KSGSSGRSTPSLSGGSGSDPAPGKLFVGGLSWQTSSDKLKEYFNMFGTVTDVLIMKDPVT 173
              ************************************************************
                                        RRM-1
Msi_PE        QRSRGFGFITFQEPCTVEKVLKVPIHTLDGKKIDPKHATPKNRPRQANKTKKIFVGGVSQ 284
Msi_PD        QRSRGFGFITFQEPCTVEKVLKVPIHTLDGKKIDPKHATPKNRPRQANKTKKIFVGGVSQ 271
Msi-PBCF      QRSRGFGFITFQEPCTVEKVLKVPIHTLDGKKIDPKHATPKNRPRQANKTKKIFVGGVSQ 300
Msi-PA        QRSRGFGFITFQEPCTVEKVLKVPIHTLDGKKIDPKHATPKNRPRQANKTKKIFVGGVSQ 272
Msi_PH        QRSRGFGFITFQEPCTVEKVLKVPIHTLDGKKIDPKHATPKNRPRQANKTKKIFVGGVSQ 233
              ************************************************************
                                                             RRM-2
Msi_PE        DTSAEEVKAYFSQFGPVEETVMLMDQQTKRHRGFGFVTFENEDVVDRVCEIHFHTIKNKK 344
Msi_PD        DTSAEEVKAYFSQFGPVEETVMLMDQQTKRHRGFGFVTFENEDVVDRVCEIHFHTIKNKK 331
Msi-PBCF      DTSAEEVKAYFSQFGPVEETVMLMDQQTKRHRGFGFVTFENEDVVDRVCEIHFHTIKNKK 360
Msi-PA        DTSAEEVKAYFSQFGPVEETVMLMDQQTKRHRGFGFVTFENEDVVDRVCEIHFHTIKNKK 332
Msi_PH        DTSAEEVKAYFSQFGPVEETVMLMDQQTKRHRGFGFVTFENEDVVDRVCEIHFHTIKNKK 293
              ************************************************************

Msi_PE        VECKKAQPKEAVTPAAQLLQKRIMLGTLGVQLPTAPGQLIGARGAGVATMNPLAMLQNPT 404
Msi_PD        VECKKAQPKEAVTPAAQLLQKRIMLGTLGVQLPTAPGQLIGARGAGVATMNPLAMLQNPT 391
Msi-PBCF      VECKKAQPKEAVTPAAQLLQKRIMLGTLGVQLPTAPGQLIGARGAGVATMNPLAMLQNPT 420
Msi-PA        VECKKAQPKEAVTPAAQLLQKRIMLGTLGVQLPTAPGQLIGARGAGVATMNPLAMLQNPT 392
Msi_PH        VECKKAQPKEAVTPAAQLLQKRIMLGTLGVQLPTAPGQLIGARGAGVATMNPLAMLQNPT 353
              ************************************************************

Msi_PE        QLLQSPAAAAAAQQAALISQNPFQVQNAAAAASIANQAGFGKLLTTYPQTALHSVRYAPY 464
Msi_PD        QLLQSPAAAAAAQQAALISQNPFQVQNAAAAASIANQAGFGKLLTTYPQTALHSVRYAPY 451
Msi-PBCF      QLLQSPAAAAAAQQAALISQNPFQVQNAAAAASIANQAGFGKLLTTYPQTALHSVRYAPY 480
Msi-PA        QLLQSPAAAAAAQQAALISQNPFQVQNAAAAASIANQAGFGKLLTTYPQTALHSVRYAPY 452
Msi_PH        QLLQSPAAAAAAQQAALISQNPFQVQNAAAAASIANQAGFGKLLTTYPQTALHSVRYAPY 413
              ************************************************************

Msi_PE        SIPASAATANAALMQAHQAQSVAAAAHHHQQQQQQQHHHQQQTHNAHVAAAQQQQQSHHN 524
Msi_PD        SIPASAATANAALMQAHQAQSVAAAAHHHQQQQQQQHHHQQQTHNAHVAAAQQQQQSHHN 511
Msi-PBCF      SIPASAATANAALMQAHQAQSVAAAAHHHQQQQQQQHHHQQQTHNAHVAAAQQQQQSHHN 540
Msi-PA        SIPASAATANAALMQAHQAQSVAAAAHHHQQQQQQQHHHQQQTHNAHVAAAQQQQQSHHN 512
Msi_PH        SIPASAATANAALMQAHQAQSVAAAAHHHQQQQQQQHHHQQQTHNAHVAAAQQQQQSHHN 473
              ************************************************************

Msi_PE        AVSNPASQAHSAAAAAALAANAANGAGAAGAHSLAAAAQQAGLMAGNPLNAAAAAAAAAA 584
Msi_PD        AVSNPASQAHSAAAAAALAANAANGAGAAGAHSLAAAAQQAGLMAGNPLNAAAAAAAAAA 571
Msi-PBCF      AVSNPASQAHSAAAAAALAANAANGAGAAGAHSLAAAAQQAGLMAGNPLNAAAAAAAAAA 600
Msi-PA        AVSNPASQAHSAAAAAALAANAANGAGAAGAHSLAAAAQQAGLMAGNPLNAAAAAAAAAA 572
Msi_PH        AVSNPASQAHSAAAAAALAANAANGAGAAGAHSLAAAAQQAGLMAGNPLNAAAAAAAAAA 533
              ************************************************************

Msi_PE        NPAAAYSNYALANVDMSSFQGVDWSTMYGMGMYV    618
Msi_PD        NPAAAYSNYALANVDMSSFQGVDWSTMYGMGMYV    605
Msi-PBCF      NPAAAYSNYALANVDMSSFQGVDWSTMYGMGMYV    634
Msi-PA        NPAAAYSNYALANVDMSSFQGVDWSTMYGMGMYV    606
Msi_PH        NPAAAYSNYALANVDMSSFQGVDWSTMYGMGMYV    567
              **********************************
```

**Supplementary Figure 2. Clustal O Multiple Sequence Alignment.** Alignment of 5 Msi protein isoform sequences showing the peptide sequence used to make the Msi antibody (2) (highlighted in yellow) and the conserved RNA-recognitions motifs (RRM-1, green; RRM-2, blue).

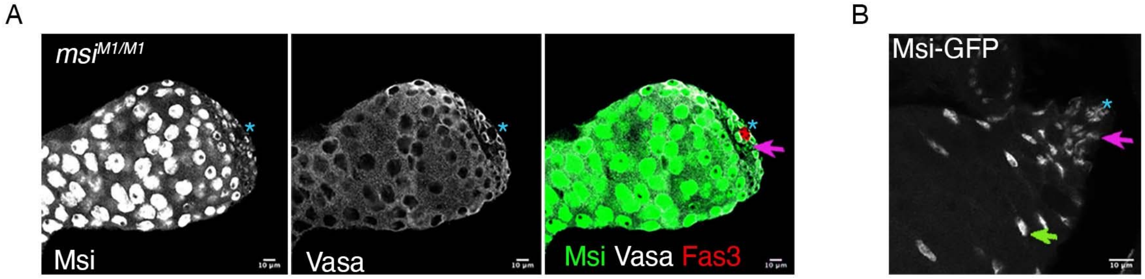


**Supplementary Figure 3. Differential expression of Msi isoforms in the *Drosophila* adult testis.** (A) Single-plane confocal micrograph showing Msi expression (green in merged panel) in an adult testis dissected from a $msi^{M1/M1}$ homozygote. Msi expression is observed in the germline and cyst progenitor cells (pink arrow) thus demonstrating that the short Msi isoforms are normally expressed in these cell-types. Blue asterix denotes the hub. (B) Single-plane confocal micrograph showing Msi-GFP expression in hub cells (blue asterix), cyst progenitor cells (pink arrow) and differentiated cyst cells (green arrow) of an adult testis.

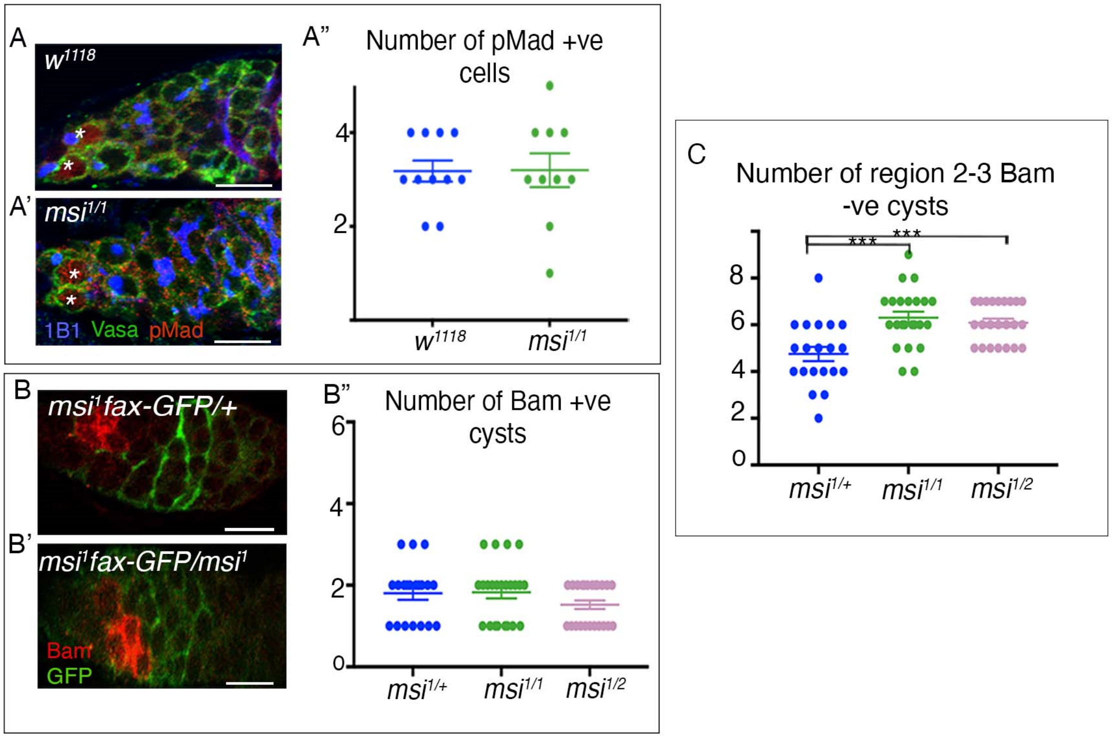


**Supplementary Figure 4. Loss of Msi function from somatic and germ cells of the ovary results in an increase in the number of germline cysts in regions 2-3 of the ovary.** (A-A') Representative single-plane confocal micrographs showing an ovary dissected from a *w1118* (A) or $msi^{1/1}$ (A') adult and labelled with antibodies to detect pMad (red), Vasa (green) and 1B1 (blue). GSCs are labelled (*). (A") Scatterplot showing the number of pMad positive cells in ovaries dissected from *w1118* or $msi^{1/1}$ adults. No significant difference between the genotypes was observed. (B-B') Representative single-plane confocal micrographs showing an ovary dissected from a $msi^{1}$*fax-GFP/+* heterozygote adult (B) or $msi^{1}$*fax-GFP/* $msi^{1}$ homozygote adult. Ovaries were labelled with antibodies to detect Bam expression (red). (B") Scatterplot depicting the average number (± SEM) of Bam-positive germline cysts in heterozygote control (blue; 1.8±.16; N=20), $msi^{1/1}$ (green; 1.82±.149; N=23) and transheterozygote $msi^{1/2}$ flies (pink; 1.52±.11; N=23). No significant difference between the genotypes was observed. (C) Scatterplot of the average (± SEM) number of Bam negative germline in region 2-3 of the germarium (e) in heterozygote control (blue; 4.75±.31; N=20), *msi*$^{1/1}$ (green; 6.304±.255; N=23) and transheterozygote *msi*$^{1/2}$ flies (pink; 6.087±.177; N=23). Welch's two-tailed t-tests show a significant increase in the number of germline cysts posterior to the region of Bam expression in both *msi*$^{1/1}$ (p=.0004) and *msi*$^{1/2}$ (p=.0007) germaria compared to control. Scale bars, 10 µm.

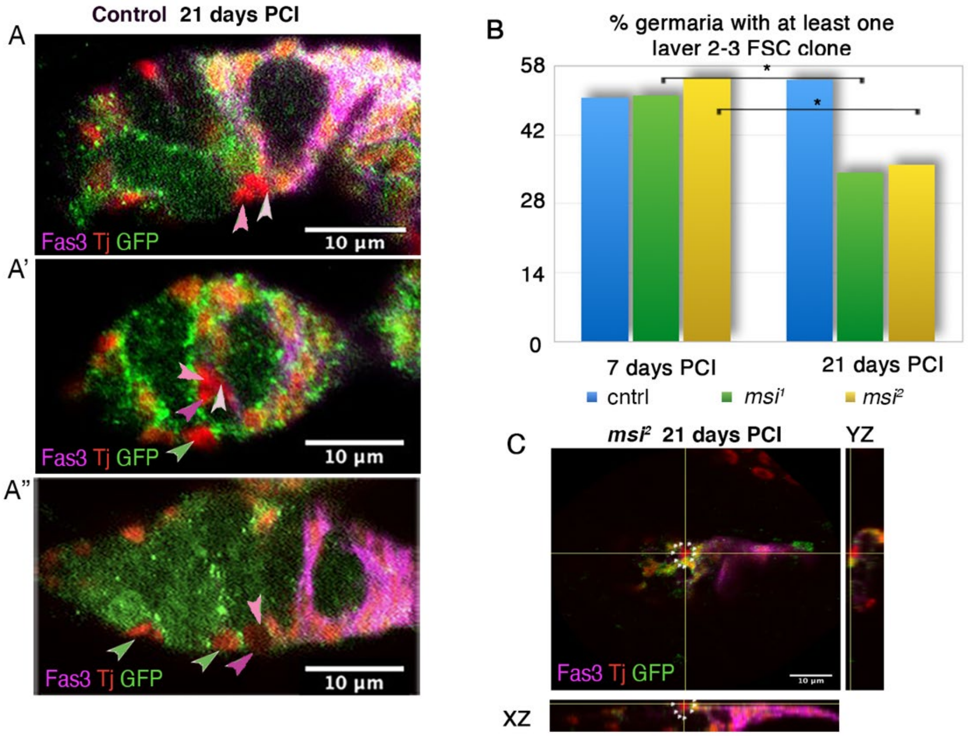


**Supplementary Figure 5. Msi loss of function results in a slight but significant decrease in proliferatively active layer 2-3 FSCs.** (A-A'') Single-plane confocal micrographs showing Traffic Jam (Tj, red) positive, GFP-negative clones generated by Flp-FRT. Layer 1 FSC clones (light pink arrowhead), layer 2 FSC clones (medium pink arrowhead) and a layer 3 FSC clone (purple arrowhead) are depicted. EC clones (green arrowheads) are also depicted. Scale bars, 10µm. (B) The percentage of germaria containing at least on negatively marked control (blue), *msi*$^1$ (green) or *msi*$^2$ (yellow) layer 2-3 FSC clone generated by Flp-FRT mediated recombination 7 and 21 days PCI. A significant reduction in the frequency of *msi*$^1$ (34.31%, N=102; Fisher's exact test p=.03) and *msi*$^2$ (36%, N=100; Fisher's exact test, p=.02) mutant FSC clones at 21 days PCI was observed.
